# The-LHON-Enigma: explaining the behaviour of Leber’s Hereditary Optic Neuropathy by the use of a simple computer model

**DOI:** 10.1101/000935

**Authors:** Ian Logan

## Abstract

Leber’s Hereditary Optic Neuropathy (LHON) appears as an ***enigmatic*** condition; affecting only certain families and often causing a severe loss of vision seemingly at random amongst family members. The first breakthrough came in 1988 with the linking of the condition to a mutation in the mitochondrial DNA (mtDNA). Now it is known that about 90% of cases are linked to 3 mutations. In this paper the *hypothesis* is suggested that a LHON mutation decreases the function of the mitochondrial enzyme, Complex I, by 50% and this alone critically endangers the survival of cells - especially the fragile cells of the optic nerves. A computer model has been written to illustrate how the *hypothesis* can produce a *natural history* for the condition of LHON that has features similar to those observed in practice; thereby successfully explaining the behaviour of this **enigmatic** condition.

## INTRODUCTION

Leber’s Hereditary Optic Neuropathy appears as an ***enigmatic*** condition; and has been so since the first cases were described in the nineteenth century by the skilled ophthalmologist Theodor Leber, 1840-1917, (Leber, 1871). He identified from amongst his patients with blindness some people who were affected by a specific condition in which the blindness was almost complete and had usually come on suddenly between the ages of 15 and 25. The condition affected men severely, but occasionally women were affected, although they often retained a degree of useful vision. However the most striking feature of the condition was that the cases of blindness appeared in families and seemingly occurred at random amongst the family members from different generations.

In the years following the initial description, the condition was given the eponymous label of Leber’s Hereditary Optic Neuropathy (LHON) and despite its rarity, the condition was well known because of its strange characteristics and its unusual inheritance. It was clear that the condition was carried down the female line, because the blindness would only appear in the children of a woman who came from a family with the condition, but would never appear in the offspring of any of the men, even if they themselves were blind. *Mendelian inheritance* struggled to make any sense of the way in which LHON was inherited as obviously the condition was not dominant and neither did it obey the rules suggestive of a recessive disease. So for many years the condition was a complete **enigma**.

> My first experience with LHON was some years ago when a young man came to see me because he was having difficulty with the sight in one of his eyes. The eye was painless, but his poor vision was starting to concern him. A local ophthalmologist was soon to diagnose LHON because of a number of typical factors: on testing it was shown that both eyes were affected, the patient’s sister also had abnormal vision, and their mother said there was an uncle, in Germany, who had lost his sight – which at the time had been put down to disease. Unfortunately, my patient went on to lose most of the vision in both eyes, but his sister was much less affected and their mother seemed totally free of any trouble. This family showed the typical presentation of LHON and many of its **enigmatic** features.

### The Link to Mitochondrial DNA

The first breakthrough in the understanding of some of the features in LHON came in the 1980’s with the realisation that inheritance was not just *Mendelian*, via genes on chromosomes, but could also come from the DNA in the mitochondria (mtDNA). Anderson and his group (1982) published the first human mtDNA sequence and this was followed by Wallace (1988) publishing a paper that linked LHON with a particular mtDNA mutation, in this case G11778A. Wallace wrote ‘A mitochondrial DNA replacement mutation was identified that correlated with this disease in multiple families.’ Since this first paper linking LHON with a mtDNA mutation there has been a veritable flood of other papers all confirming the causal link. This high interest in publishing papers showing the results of new studies has continued to the present. Over the years researchers have shown that the 3 main LHON mutations in the mtDNA are G3460A, G11778A & T14484C, and they account for about 90% of cases, whilst the other 10% of cases are caused by the many less common mtDNA mutations.

### The Spectrum of LHON Disease

It has also become apparent that LHON is not a single condition but a syndrome; with, in most instances, a person showing only a unilateral or a bilateral visual deficit. However in some people there are obvious muscular, cardiac and neurological symptoms and signs; and on theoretical grounds this would not be unexpected as mitochondria occur in all the cells of the body. LHON appearing with some extra features has been described by using the term ‘LHON PLUS’, or similar, (Nikoskelainen, 1995). But in people whose condition fits the description of ‘LHON PLUS’, there will still be an underlying mitochondrial mutation; and it is to be expected that the optic neuropathy will feature prominently.

Also, there are instances when the condition of LHON appears to be **mild**. In people with this form of LHON it maybe that the mitochondrial mutation in the person is heteroplasmic for a common LHON mutation, or it may just be the person is a member of a family that appears to have a low incidence of LHON symptoms.

On the other hand, there are also instances when LHON appears to be **severe**, producing sudden and severe visual deficit in almost every family member, either because of the presence of a particularly active mitochondrial mutation, or just because they come from a family which has a high incidence of problems.

When a mutation causes very severe problems the condition can take the form of ‘subacute necrotizing encephalomyelopathy’, or Leigh’s disease; named after a British psychiatrist Dr. Denis Archibald Leigh, 1915-1998. However, in practice the clinical syndrome can be caused by various autosomal mutations (Marin, 2012), as well as mtDNA mutations, such as T12706C (Taylor, 2002).

LHON, therefore, has a spectrum of presentations, from the **mild** form where there is often no subjective loss of a visual deficit, and other relatives may only have slight loss of vision; though the **average** form where a person will usually have a slight degree of visual deficit, perhaps only noticeable on testing (Sacai, 2010) and sporadic family members will have a marked visual deficit; to the **severe** form, where severe optic atrophy affects almost everyone, as well as, on occasions, causing other serious disablements.

> In the computer model the different forms of LHON are divided into the three groups **mild**, **average** and **severe**.

### Factors Adversely Affecting LHON

A variety of factors have been suggested that may precipitate a sudden deterioration in visual acuity; and the most widely accepted damaging factors are smoking and heavy alcohol intake. (Kirkman, 2009).

There are also several interesting reports of visual acuity worsening after exposure to polycyclic aromatic hydrocarbons (Rufa, 2005) and n-hexane (Carelli, 2007); and, in the laboratory, 2,5-hexanedione (Ghelli, 2009) and rotenone (Santos, 2006; Marella, 2010) have also been shown to influence cell cultures.

There is also evidence that a LHON-LIKE optic neuropathy may be caused by malnutrition, the abuse of alcohol and smoking, even in persons who do not have any known DNA mutation. The optic neuropathy in these cases can appear clinically similar to LHON, but there will not be the characteristic maternal inheritance. The best example, in this respect, is the outbreak in the 1990’s of *Cuban Epidemic Neuropathy* in which as many as 50,000 Cubans were affected to a greater or lesser degree. There were a number of papers produced during the epidemic, or shortly afterwards (Johns, 1994; Sadun, 1994; Sadun, 1994-1995; Sadun, 1998; Santiesteban-Freixas, 1999); and more recently further papers have appeared (Santiesteban-Freixas, 2010; Mills, 2011) which look at the epidemic with our present knowledge.

> In the computer model the factors of exposure to toxins, smoking, alcohol intake and malnutrition are considered together as one parameter which can be set as **none**, **mild**, **moderate**, or **severe**.

### The Importance of Family History

One of the features of LHON is that the disease appears to vary from one family to another with some families apparently doing better than others, although the families may share the same mitochondrial mutation. The reasons for these differences are not clear, but they presumably arise as a combination of both the autosomal background (other mutations that affect mitochondrial function) and environmental factors.

> In the computer model a person’s family history is considered as **poor**, **average**, or **good**.

### The Difference between the Sexes

Much has been made of the way in which LHON commonly affects the males of a family much more frequently than the females; and this difference was noted by Theodor Leber (1871) in his original study. Indeed, in some families it appears that LHON is a disease of the men only, as all the women appear to be free of the signs and symptoms of the condition.

However, the question of whether or not females have a protection against LHON is debatable. Sacia (2010) in a study shows just how close their symptom-free women are to being affected. Whilst, Giordano (2011) working with cell cultures has shown that oestrogens may be beneficial to mitochondrial function. So whether females have **protection**, or males are **adversely affected** by toxins, smoking and alcohol intake is undecided.

> In the computer model a person’s sex is not taken to be an independent variable.

### Age of Onset of Symptoms

LHON rarely affects children, but it can occur (Barboni, 2006). However, symptoms most commonly develop in the late teenage years, rather than at any other age. There has never been a good explanation of why this should happen. But recent evidence does suggest that children have more mtDNA molecules, and presumably more mitochondria, than older persons and this may turn out to be the factor determining why children do better than adults (Chu, 2012).

> In the computer model there is a low *risk* of developing loss of vision before age 15.

### The Mitochondrial Enzyme Complex I (NADH-ubiquinone oxidoreductase)

The mitochondrial mutations that are associated with LHON are all to be found in the nucleotide bases of the genes for peptides that form subunits of the mitochondrial enzyme **Complex I**, or to use the more descriptive name, **NADH-ubiquinone oxidoreductase**.

In respect of the common LHON mutations:

**G11778A** - occurs in the gene for the peptide MT-ND4 (NAD4) and changes the 340^th^ amino acid from an *Arginine* to a *Histidine*.
This is a subtle change as the amino acids are similar to each other and both carry an electrical charge which makes them important in making links within proteins. However, a *Histidine* has a weaker positive charge and is physically slightly shorter than an *Arginine*, and this appears enough to disrupt the function of the peptide.
**T14484C** - occurs in the gene for the peptide MT-ND6 (NAD6) and changes the 64^th^ amino acid from a *Methionine* to a *Valine*. Again this change is subtle as the amino acids are both neutral in charge. However, *Methionine* contains a sulphur atom and is physically larger than a *Valine*, and these differences appear to be enough to effect a change in function.
**G3460A** - occurs in the gene for the peptide MT-ND1 (NAD1) and changes the 52^nd^ amino acid from an *Alanine* to a *Threonine*. Again the change is subtle, but a *Threonine* is physically larger and this appears to affect the function of the peptide.

And, making a similar analysis for mutations that cause Leigh’s disease:

**T12706C** – occurs in the gene for the peptide MT-ND5 (NAD5) and changes the 124^th^ amino acid from a *Phenylalanine* to a *Leucine*. Once again, this is only a subtle change as the amino acids are very similar, but *Phenylalanine* is larger. (Taylor, 2002)
**G14600A** – occurs in the gene for the peptide MT-ND6 (NAD6) and changes the 25^th^ amino acid from a *Proline* to a *Leucine*. This is a more marked change as a *Proline* is considerably larger than a *Leucine*. For the case report in respect of this mutation see Malfatti (2007) and for a mouse model (Lin, 2012).

Complex I is a large enzyme and is described as being the first enzyme in the mitochondrial electron transfer chain. Molecules of this enzyme are found associated with the inner membrane of the mitochondrion. Each molecule of enzyme appears to be L-shaped with a hydrophilic part positioned internally to the membrane, and a trans-membranous part which contains channels through which ions can be pumped.

In the simplest of terms the enzyme uses the energy released in the reaction

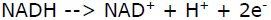

and pumps the H^+^ ions from the mitochondrial matrix, through the inner membrane of the mitochondrion into the space beyond; and this generates an electrical potential that later in the electron transfer chain can be used to produce ATP from ADP.

Each molecule of Complex I is made by assembling about 45 peptide subunits; 7 of which are encoded by the mtDNA. These peptides are MT-ND1, MT-ND2, MT-ND-3, MT-ND4, MT-ND4L, MT-ND5 & MT-ND6; and all of these peptides are present in the transmembranous part of the enzyme (Roberts, 2012).

Recent work by Dröse (2011) suggests that there are actually 2 separate ion pumps; a proximal pump which uses the transmembranous channels of the MT-ND1, MT-ND2, MT-ND3, MT-ND4L and MT-ND6 peptides, and a distal pump which uses the channels of just the MT-ND4 and MT-ND5 peptides.

Also it is suggested that each pump is of equal power; which leads to the conclusion that 50% of a mitochondrion ion pumping power is reliant on the proper functioning of the MT-ND1, MT-ND2, MT-ND3, MT-ND4L and MT-ND6 peptides, whilst the other 50% is reliant on the MT-ND4 and ND-ND5 peptides.

Figure 1, adapted from Dröse (2011) gives a suggested structure for Complex I.

> The computer model uses the suggestions from Dröse (2011) and considers that the presence of a LHON mutation decreases mitochondrial function by 50%.

**Figure 1.**
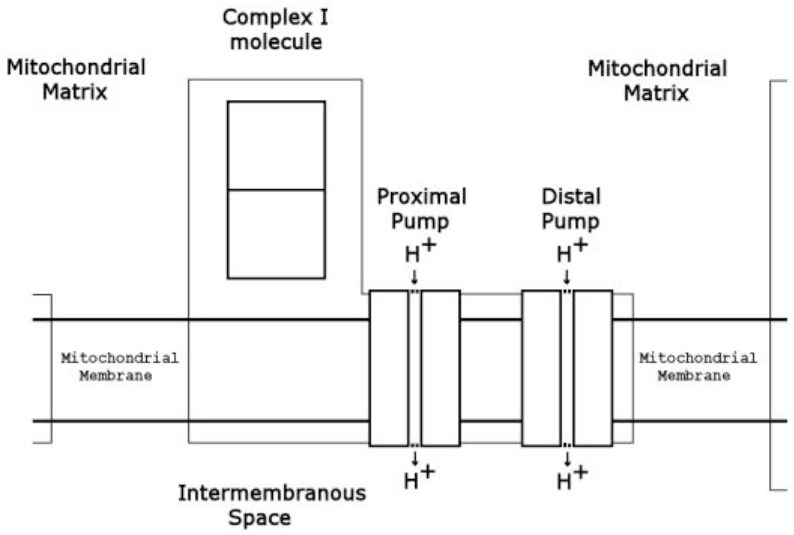
Suggested Structure of Complex I (shaded area). Each molecule is L-shaped and has two proton pumps. H^+^ ions are pumped from the Mitochondrial matrix through the Mitochondrial membrane into the Intermembranous space. Figure adapted from Dröse (2011).

### Overall Purpose

This paper offers a *hypothesis* and a computer model to explain the **enigmatic** behaviour of LHON. The work uses the suggestions contained in the recent paper by Dröse (2011) to show how LHON mutations decrease the functioning of the enzyme Complex I.

The paper also gives a scientific basis to the the condition of LHON which may lead to therapy that prevents the onset of blindness in persons with mtDNA mutations. In the discussion the characteristic features of LHON will be considered one by one, and the *hypothesis* tested against each in turn. It is not possible to say that all the features of LHON can now be explained but the *hypothesis* and the computer model do appear to be useful.

## METHODS

This paper and the accompanying computer model are the result of a project undertaken to produce a guide to the genetics of LHON for publication on the Internet. At the start of the project, a search was made on GenBank for published mtDNA sequences with common LHON mutations. This search found 46 mtDNA sequences with G11778A, 33 with T14484C, and 19 with G3460A. On GenBank sequences with these mutations are over-represented, perhaps by a factor of 3, because of the scientific interest in the condition; but the rough proportions of 3:2:1 for the common mutations would appear to be reasonable.

Next, a search was made for scientific papers dealing with LHON for abstracts. Many full reports were also collected. Of special interest is the original report of LHON cases by Theodor Leber (1871); and although printed copies are extremely rare, a copy of his original work is now available on the Internet at: http://link.springer.com/journal/417/17/2/page/1

From the collection, two recent papers were noted to be particularly importance in respect of LHON; although in neither case were the authors involved with the clinical care of persons with LHON.

The paper by Dröse (2011) titled ‘Functional dissection of the proton pumping modules of mitochondrial complex I’, describes how complex I has two proton pumps of equal power; a proximal proton pump constructed from some mtDNA peptides and a distal pump constructed from other peptides.

And, the paper from Lin, et al. (2012) titled ‘Mouse mtDNA mutant model of Leber hereditary optic neuropathy’, describes how ‘chronic oxidative stress’ causes the death of the cells of the optic nerve – effectively one cell at a time as each gets overwhelmed by ‘reactive oxygen species’ (ROS).

This latter paper is also important because it describes ‘abnormal mitochondrial morpholgy and proliferation’ being observed in the cells of the optic nerves as they try to withstand the stresses placed upon them.

The ideas in these papers were then used to develop a *hypothesis* to explain the behaviour of LHON. Subsequently a computer model using this *hypothesis* was written so as to demonstrate in a simple way how recent advances in the understanding of the scientific basis of the condition have helped to explain its **enigmatic** features.

## DISCUSSION

### A *HYPOTHESIS* TO EXPLAIN LHON

The condition of LHON usually presents as a sudden loss of visual acuity affecting firstly one eye and then the other. The condition appears sporadically in family members with the males being more likely to be affected and more severely affected that the female member of the family. The peak age for the onset of optic nerve failure appears to be between the ages 15 to 25; but the condition does occasionally occur in children. In persons over the age of 50 it is unusual to have problems for the first time.

In most families with LHON, a mitochondrial mutation is to be found in the mtDNA of the affected persons; and the 3 *common* mutations, G3460A, G11778A, T14484C account for over 90% of the cases. Fortunately, in most affected people no further disabilities are to be found other than the loss of visual acuity.

But LHON clearly has a spectrum of presentations, with some people having the appearances of a *mild* condition with only partial visual loss, whilst in other people the condition is *severe*, with the loss of vision being just one of several disabilities.

So how might this **enigmatic** behaviour of LHON be explained?

In the author’s opinion the 2 papers by Dröse (2011) and Lin (2012) give some good insights into the scientific basis of LHON; and these insights do provide the basis for the *hypothesis* that:

> *LHON is a condition in which the functioning of the mitochondrial enzyme Complex I is reduced by 50% and this alone critically endangers the survival of the cells of the body*.

So to see how this *hypothesis* might work in practice consideration must be given to the dynamics of energy *demand* and *supply* within the cells of the body. The following explanation is purposely kept very simple and general; and the computer model written for this study uses many of the points discussed in its program.

### USING THE *HYPOTHESIS* TO EXPLAIN LHON

#### The *supply* of energy

**When the mtDNA is *normal***: for each cell of the body it is possible to consider that the *supply* of energy is largely the result of the ***normal*** functioning of the enzyme Complex I. The pumping of H^+^ ions through the inner membrane of the mitochondrion is not impaired and there is a good potential difference that can in turn be used to convert ADP to ATP (Gouspillou, 2011).

The *supply* of energy is therefore proportional to the functional ability of the enzyme Complex I; and when the 2 proton pumps of the enzyme are both fully functional the *supply* of energy will be quite sufficient to maintain the survival of the each mitochondrion, and in turn the complete cell. This includes the rather fragile nerve cells of the optic nerves where *normal* mtDNA allows for an adequate supply of energy.

However, there are some other points that should be considered:

Firstly, the functional ability of enzyme Complex I can become impaired by other factors such as chemicals, alcohol, smoking and malnutrition; and as a result the cells of the optic nerve may not survive when adversely affected in this way, notwithstanding the presence of *normal* mtDNA.

The second point to consider is the important fact that the *supply* of energy in a mitochondrion, and hence the whole cell, will vary with time. It is a feature of all biological processes that there is variation with no process being absolutely constant. So, the *supply* of energy in a cell will vary; and this variation can be considered to follow a normal distribution. There will therefore be occasions when the cell’s capacity to *supply* energy will be appreciably lower than at other times. Barboni (2012) uses the term ‘waxing and waning’ to describe this phenomenon.

**When a LHON mutation is present**: the functional ability of the complex I molecules is markedly reduced to about the 50% point, as only one proton pump will be available in each molecule as opposed to the normal two pumps. This places the fragile cells of the optic nerves at considerable risk as the mitochondria within these cells have a much reduced capacity to *supply* energy.

> In the computer model a cell’s capacity to *supply* energy follows a normal distribution; resulting in periods when there is insufficient supply to maintain the survival of the cells of the optic nerve.

#### The *demand* for energy

As with the *supply* of energy within a cell, the *demand* for energy fluctuates (Calmettes, 2008). It is unclear what exactly might cause the changes, but in simple terms it can be considered that a high ADP level, and a corresponding low ATP level, within a cell places a *demand* on the mitochondria to increase their production of energy.

Again, the *demand* for energy can be considered to follow a normal distribution with some periods of low *demand*, an average *demand* for most of the time, and some occasions when there is a high *demand*.

#### The result of high *demand* and low *supply*

So what happens when the *demand* for energy is high and the cell finds that it needs to raise the *supply* of energy.

In this situation it would appear that there is a stimulus to the manufacture of new mitochondria, together with new complex I molecules, in an attempt to rectify the shortage, a process termed mitochondrial biogenesis. However, this process, which appears to involve for the most part the fission of existing mitochondria, will clearly takes some time; and in the meanwhile the cell will be at considerable risk.

The recent paper from Lin (2011) discusses the finding of mitochondrial proliferation and mitochondria with abnormal morphology in cells that are struggling to survive under these circumstances – with the implication that there is a high *demand* for energy that the cell is unable to satisfy; and the normal processes within a cell become disrupted.

Figure 2 shows the way in which the normal distribution curves of the *demand* for energy and the *supply* do not overlap when the mtDNA is normal (and there are no toxic factors to consider). However, figure 3 shows that in the presence of a LHON mutation, there is an overlap of the normal distribution curves; which may lead to the pattern of cell death that is typical of the condition of LHON.

**Figure 2.**
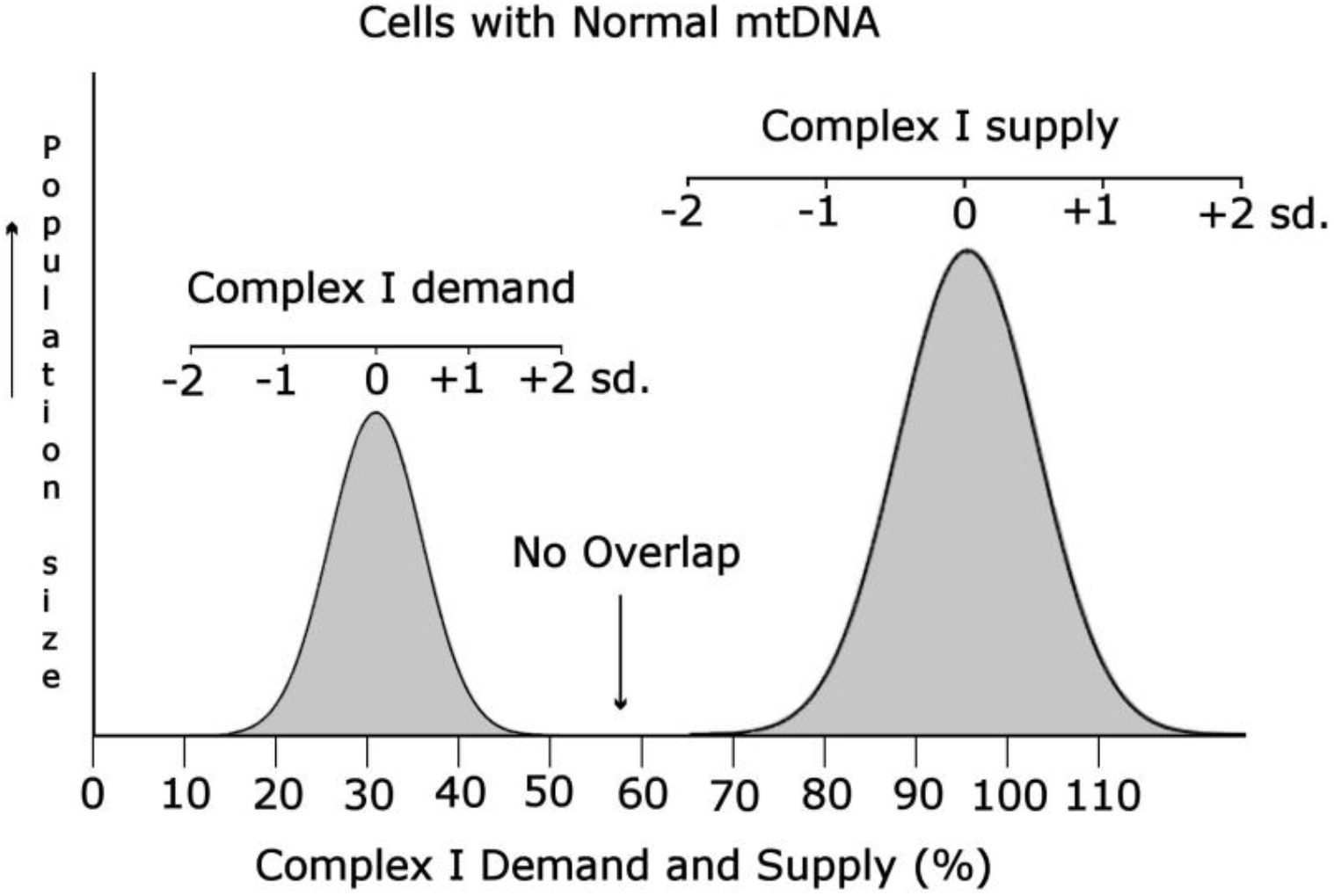
shows how the *Demand* for energy and the *Supply* follow a normal distribution; and in cells with Normal mtDNA these distributions can be considered not to overlap. I.e. the *Demand* can always be satisfied.

**Figure 3.**
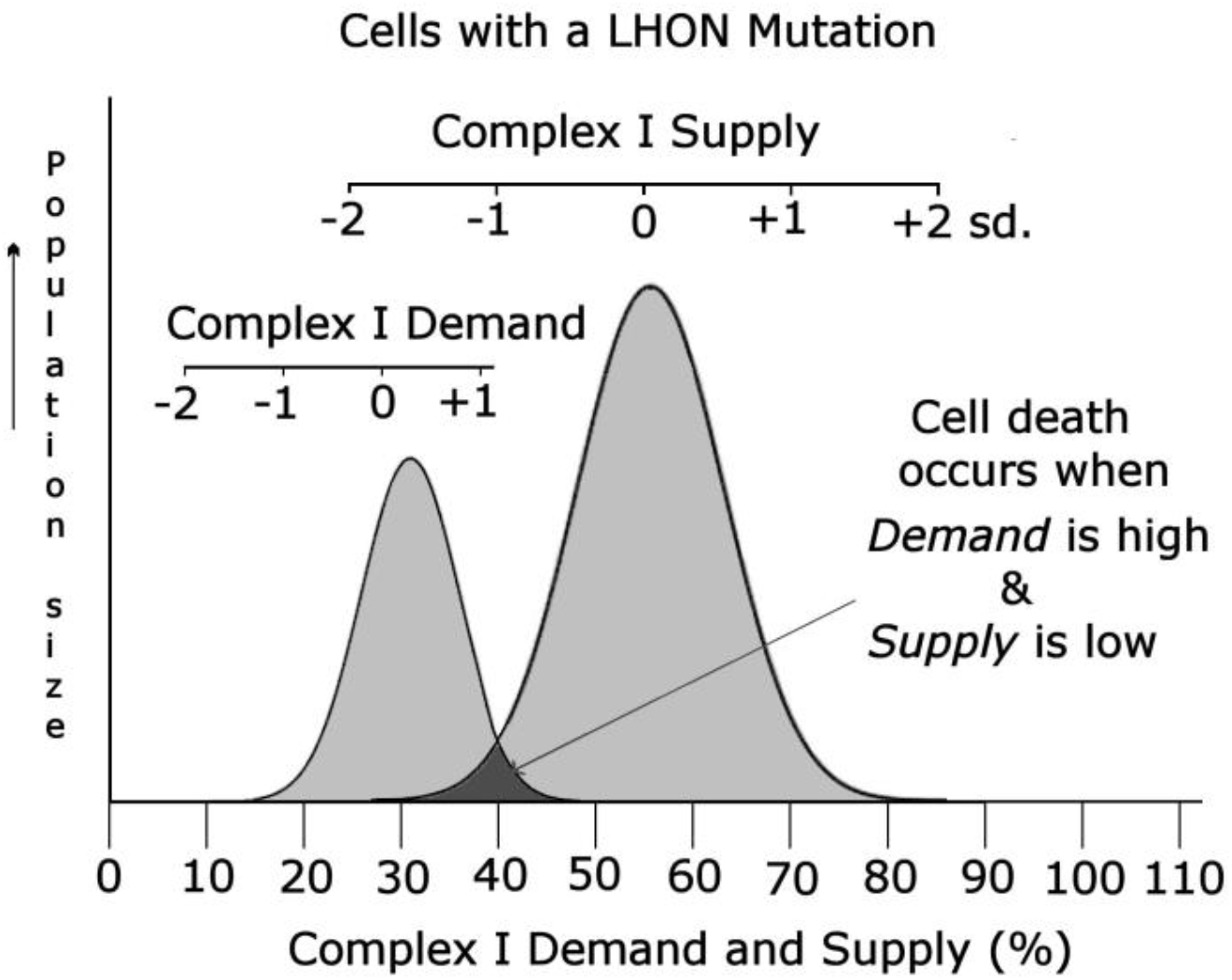
shows the situation in cells which have a LHON mutation in their mtDNA. Here the capacity of the cells to supply energy is compromised; and when the *demand* for energy is high and the *Supply* of energy is low death of the fragile cells in the optic nerves may occur.

### The Modelling Process

As part of the present study a computer model was produced to illustrate how the *hypothesis* presented above can be used to reproduce a *natural history* for the condition of LHON that has features similar to those observed in practice.

LHON has many **enigmatic** features, and the computer model attempts to show how the *hypothesis* resolves the problems by following the survival of the cells of the optic nerves throughout a person’s life. The model allows for various parameters to be changed and *the person re-born under different circumstances* so that the effect of the revised parameters can be observed in a simple way.

For the computer model a series of assumptions are made:

- the thousands of cells of the optic nerves can be represented by the left optic nerve having 128 cells and the right optic nerve another 128 cells. The decision to model the fate of just 256 cells was fairly arbitrary, but this number of cells can be displayed in a simple manner.
- the model shows the survival, or the death, of the cells of the optic nerve over the lifetime of a person; and in the model the lifetime is considered in 1000 periods of time, each of which corresponds to a month.
- the LHON mutation can be considered as being **mild**, **common** or **severe**.

**Mild**: representing heteroplasmy, or the presence of a LHON mutation of low activity; when serious loss of vision in a lifetime is uncommon.
**Common**: representing the presence of a common LHON mutation; when marked loss of visual acuity appears sporadically amongst the family members.
And **Severe**: indicating the mutation can cause Leigh’s disease; with the optic neuropathy being just one of many problems.
- the effect of toxins, smoking, alcohol intake and malnutrition can be considered as a single parameter. The settings can be:

**None**: there is no increased risk.
**Mild**: there is some increase in risk; such that a person with a common LHON mutation will suffer some loss of visual acuity.
**Moderate**: there is a significant increase in risk; so that a person with a common LHON mutation will have a considerable loss of visual acuity.
**Severe**: there is a substantially increased risk and even person with normal mtDNA are likely to have a loss of visual acuity.
- the effect of family history. This parameter alters the sensitivity of the whole model. The settings can be:

**Poor**: indicative of a family where LHON symptoms are frequent.
**Average**: the typical setting for a family with a common LHON mutation.
**Good**: indicative of a family where LHON symptoms are uncommon
- the loss of visual acuity.

In the model a simple count is made of the numbers of *live* and *dead* cells in the optic nerves and this is expressed as a percentage. When all the cells are *live* the loss of visual acuity will be 0%, and 100% when all the cells are *dead*.

### The algorithm used in the computer model

In the program a *personal-risk* value is given to the person and is taken to be the arithmetic sum of values representing the person’s age, their mtDNA status, and the exposure to toxins. A *cellular-risk* value is also given to each of the cells of the optic nerves. These values are ordered such that cells from the core of the optic nerves have lower values than the presumed more resistant cells of the outer parts of the optic nerves to which are given higher values.

Then, for each time slice, the *personal-risk* value is randomised following the rules for a normal distribution. The new value is then compared against each of the *cellular-risk* values for those cells that are still alive. If the *personal-risk* value exceeds a *cellular-risk* value then that cell will also die.

Various randomising operations are performed on the initial data to ensure that different patterns of cell death are produced on each new *run* of the program. For example, whether it is the left eye that is to be affected predominantly, or the right eye, is determined at random.

The overall sensitivity of the program is altered by the family history setting. A **poor** family history places the cells at greater risk by lowering the *cellular-risk* values, whereas a **good** family history raises the values.

### Example displays from the computer model

The computer model produces different displays depending on the **Selected Options** chosen for the *run*.

As a demonstration of what will be produced by the computer model 4 representative displays are given here.

Display (a) – shows the defaults for the Selected Options; and the display shows all the cells of the optic nerves are *live*.

Display (b) - illustrates a *run* for a person with a *Common LHON mutation*, but without any *toxic factors*. In this instance the individual has reached age 75 and has an 11% visual deficit, affecting more one eye than the other.

Display (c) - shows a run for an individual who carries a common LHON mutation, but who also has mild toxic factors; and here, even in young adulthood there is a marked visual deficit affecting both eyes.

Display (d) - illustrates how severe toxicity from chemicals, smoking, alcohol intake or malnutrition (such as seen in *Cuban Epidemic Neuropathy*) might cause a significant loss of vision even in a person with normal mtDNA.

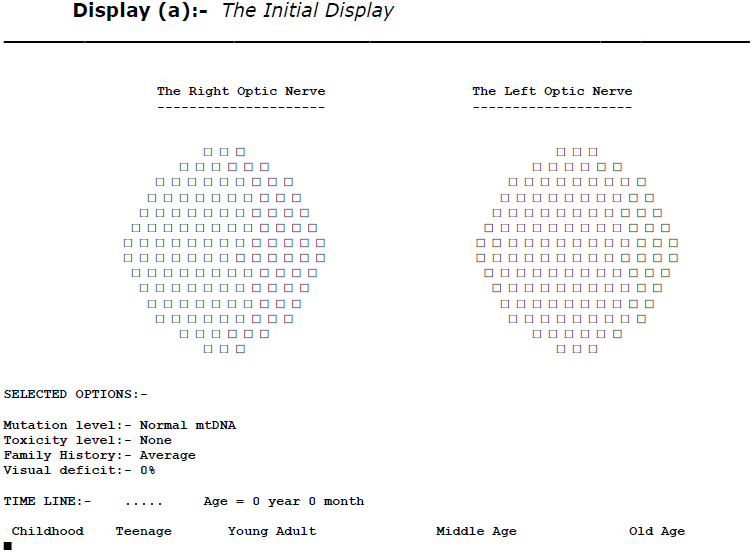
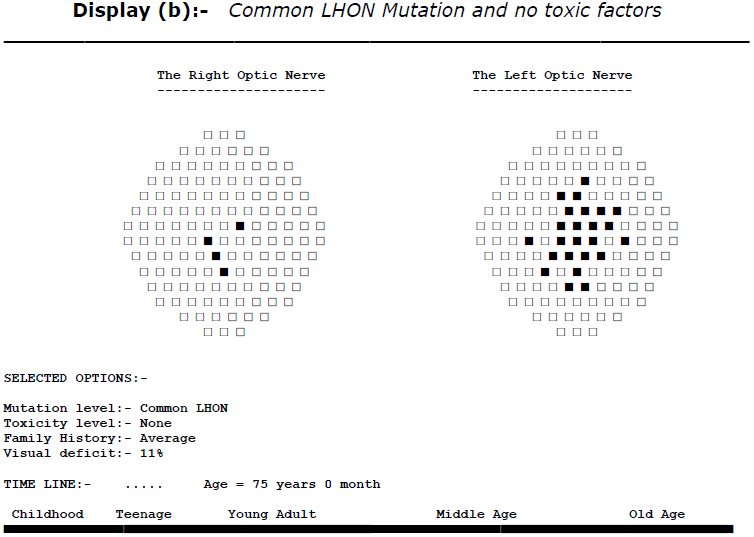
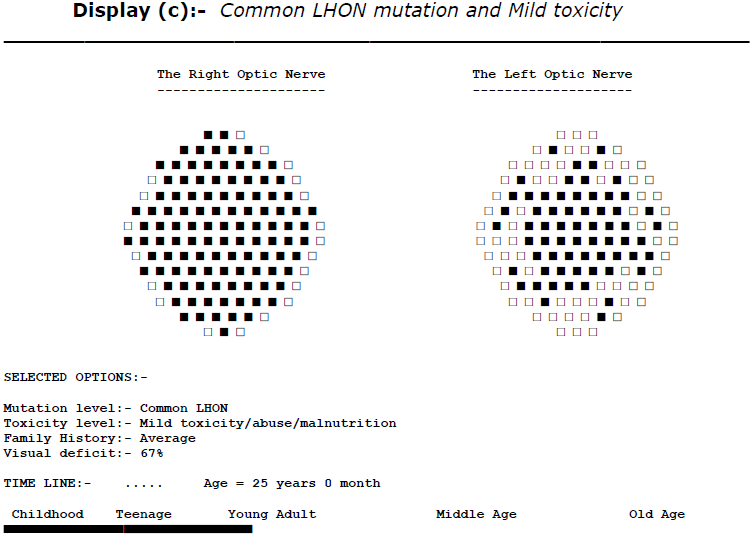
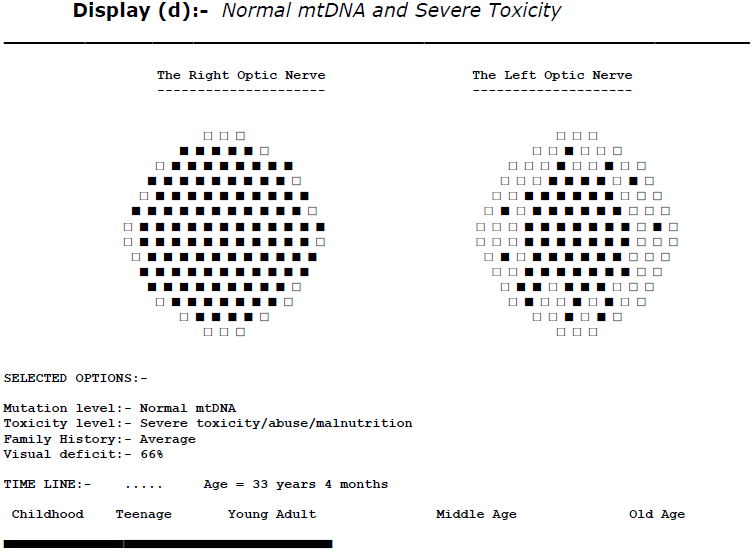

### The enigmatic features of LHON

The computer model described in this study is not a **simulation**, as it does not contain data collected from actual cases of LHON, but rather it is a **predictive model** using simple rules based on a scientific understanding of the condition.

For a person with normal mtDNA the model predicts:

> - the cells of the optic nerves will stay alive, except when there are **severe** toxic factors (as may have occurred with *Cuban Epidemic Neuropathy*).

However, for a person with a LHON mutation in their mtDNA the model predicts:

> - there is a variable risk of suffering a visual deficit; and this risk will be increased as other factors, such as exposure to toxins, alcohol intake, smoking, and an adverse family history are taken into account.

So does the computer model explain the **enigmatic** features of LHON?

The model uses the *hypothesis* that a LHON mutation decreases the ability of a cell to *supply* energy by about 50%, and as a result it is possible to explain that:

> LHON is familial – and depends on the presence of a mtDNA mutation carried down the maternal line.
>
> The loss of vision is variable – and depends on the person’s age, the mtDNA mutation, the presence of toxic factors and the family history.

To this extent the computer model gives a good explanation of how a very simple *hypothesis* can be used to show how just small variations between the inheritance of one person, together with the habits of that person, in respect of exposure to toxins, alcohol intake and smoking, may alter the extent of any visual loss suffered by that person in their lifetime; and in this way the computer model does give an explanation of the **enigmatic** features of LHON.

However, the computer model fails to explain all the features of LHON; as the model *assumes* rather than *explains* why LHON disease usually affects the males in a family more than the females, and why there is usually an absence of the condition in childhood.

### Conclusions

Leber’s Hereditary Optic Neuropathy appears to be an ***enigmatic*** condition and the causes of the condition have been puzzling researchers for many years. For example Vilkki (1991) wrote over 20 years ago ‘the variation in the clinical expression of the disease among members of the same family has remained unexplained’ and very recently Hudson (2011) wrote ‘[LHON] has an unexplained variable penetrance and pathology …… the mechanisms leading to neurodegeneration remain unclear’.

In this paper a *hypothesis* is put forward to explain the **enigma** of LHON. The *hypothesis* uses the simple suggestion that a LHON mutation can be considered to reduce the functioning of the enzyme, Complex I, by 50%; which in its turn critically endangers the survival of the cells of the optic nerves.

The computer model written for this study provides a way for seeing how the *hypothesis* might work in a variety of circumstances, such as at different ages, with a variety of different LHON mutations, and when toxic factors are also present.

The *hypothesis* and the computer model cannot be expected to answer all the **enigmatic** features of LHON, but they do appear to go some way towards increasing our understanding of this condition.

Finally, the *hypothesis* and computer model suggest that LHON symptoms come about because of a decrease in the *supply* on intracellular energy; and therefore therapy aimed at generally increasing this *supply* may be successful in preventing symptoms. At the present time it would appear we do not know how to stimulate the production of new mitochondria and Complex I molecules, the process of mitochondrial biogenesis, but learning how to do so may the key to preventing the symptoms of LHON and possibly many other conditions.

## Additional Material

Additional file: the-lhon-enigma.txt (35 K) the source file for the computer model.

## Internet resource

The-LHON-Enigma computer model is available at: www.ianlogan.co.uk/lhon/the-lhon-enigma.htm

